# ChIPWig: A Random Access-Enabling Lossless and Lossy Compression Method for ChIP-seq Data

**DOI:** 10.1101/127464

**Authors:** Vida Ravanmehr, Minji Kim, Zhiying Wang, Olgica Milenković

## Abstract

**Motivation:** The past decade has witnessed a rapid development of data acquisition technologies that enable integrative genomic and proteomic analysis. One such technology is chromatin immunoprecipitation sequencing (ChIP-seq), developed for analyzing interactions between proteins and DNA via next-generation sequencing technologies. As ChIP-seq experiments are inexpensive and time-efficient, massive datasets from this domain have been acquired, introducing significant storage and maintenance challenges. To address the resulting Big Data problems, we propose a state-of-the-art lossless and lossy compression framework specifically designed for ChIP-seq Wig data, termed ChIPWig. Wig is a standard file format, which in this setting contains relevant read density information crucial for visualization and downstream processing. ChIPWig may be executed in two different modes: lossless and lossy. Lossless ChIPWig compression allows for random access and fast queries in the file through careful variable-length block-wise encoding. ChIPWig also stores the summary statistics of each block needed for guided access. Lossy ChIPWig, in contrast, performs quantization of the read density values before feeding them into the lossless ChIPWig compressor. Nonuniform lossy quantization leads to further reductions in the file size, while maintaining the same accuracy of the ChIP-seq peak calling and motif discovery pipeline based on the NarrowPeaks method tailor-made for Wig files. The compressors are designed using new statistical modeling approaches coupled with delta and arithmetic encoding.

**Results:** We tested the ChIPWig compressor on a number of ChIP-seq datasets generated by the ENCODE project. Lossless ChIPWig reduces the file sizes to merely 6% of the original, and offers an average 6-fold compression rate improvement compared to bigWig. The running times for compression and decompression are comparable to those of bigWig. The compression and decompression speed rates are of the order of 0.2 MB/sec using general purpose computers. ChIPWig with random access only slightly degrades the performance and running time when compared to the standard mode. In the lossy mode, the average file sizes reduce by 2-fold compared to the lossless mode. Most importantly, near-optimal nonuniform quantization with respect to mean-square distortion does not affect peak calling and motif discovery results on the data tested.

**Availability and Implementation:** Source code and binaries freely available for download at https://github.com/vidarmehr/ChIPWig

**Supplementary information:** Is available on bioRxiv.

## 1 Introduction

Next-generation sequencing (NGS) platforms are powerful data acquisition technologies that lead to invaluable information regarding genomes, transcriptomes or epigenomes of an organism. Due to the low cost of NGS technologies such as Illumina (Illumina (2014b)), one can now sequence a human genome in less than 15 hours at the cost of $1000 (Illumina (2014a)). Consequently, the number of sequencing projects and accompanying databases has experienced a tremendous growth. In the emerging data repositories, ChIP-seq files often compromise a significant fraction of the stored information. ChIP-seq is an efficient technique to analyze the interactions between protein and DNA by combining Chromatin immunoprecipitation and NGS methods. ChIP-seq experiments have been widely performed by numerous projects, including the ENCODE project (Consortium *et al*. (2004)) hosted at the UCSC Genome Browser, various National Institute of Health (NIH) programs, the Roadmap Epigenomics (Bernstein *et al*. (2010)) and the Cistrome Project (Liu *et al*. (2011)). Methods for downstream analysis of ChIP-seq data have also been reported in a vast volume of the bioinformatics literature, and they include peak calling and motif finding (Bailey *et al*. (2013); Kuan *et al*. (2011); Nakato and Shirahige (2016); Machanick and Bailey (2011)).

ChIP-seq experiments generate raw reads in FASTQ format that require large storage space and are unsuitable for visualization. Consequently, most FASTQ files are accompanied by Wig files that provide summary information contained in the reads that needs to be visualized, such as coverage, read density or expression. The sizes of ChIP-seq files in Bed and Wig format, available from ENCODE, Gene Expression Omnibus, or Galaxy repositories, often exceed 1 GB. Given that they are accessed and downloaded with high frequency, it becomes hard to avoid storage and communication bottlenecks. To enable efficient maintenance and organization of these and other genomic databases, specialized compression methods have been developed that can significantly reduce the size of the sequencing datasets (Cao *et al*. (2007); Tabus and Korodi (2008); Pinho *et al*. (2011); Kent *et al*. (2010); Hoang and Sung (2014)). For Wig files, two of the most frequently used compression techniques are bigWig and bigBed (Kent et al. (2010)). BigWig essentially uses classical gzip compression and leads to moderate reductions in file sizes. A more efficient compression method for RNA-seq and ChIP-seq Wig files, termed cWig (Hoang and Sung (2014)), was recently proposed, but appears to require special compilers and is currently not publicly available.

Recently, the authors proposed a compression method for RNA-seq Wig files termed smallWig (Wang *et al*. (2016)) that represents the state-of-the-art method in the field, offering significantly improved compression rates when compared to bigWig, cWig and gzip. SmallWig enables fast queries from the compressed files in addition to access to summary statistics features paralleling those offered by the UCSC Genome Browser. At the core of the smallWig algorithm are two separate encoding pipelines for location and expression tracks via differential and run-length encoding, and accompanying arithmetic compaction. Although smallWig offers a large gain in terms of compression rate and compression/decompression time for RNA-seq Wig files, it is not suitable for use on ChIP-seq data due to the different statistical properties of the data tracks. In RNA-seq files, the first column of the data lines represents chromosome locations that appear as consecutive positive integers, while in ChIP-seq Wig files, the chromosome locations in the first column are positive integers that are not necessarily consecutive. In ChIP-seq files, the average read densities have fairly high variances both globally and locally; by contrast, in RNA-seq Wig files, the expression values appear to be locally smooth. As a result, the smoothness feature of RNA-seq expression tracks makes them suitable candidates for run-length encoding, which may not be adequate for ChIP-seq data. Most importantly, smallWig is only designed to operate in a lossless mode, which may not be needed for highly noisy data used for visualization such as ChIP-seq Wig data. ChIP-seq data is standardly processed through sequential peak calling followed by motif discovery, which makes it possible to exactly characterize the effects of lossy data quantization on the performance of two these inference algorithms. Thus, in many applications it may be desirable to perform lossy compression of ChIP-seq data, as it significantly reduces file sizes while preserving information relevant for downstream processing.

We introduce a compression technique for ChIP-seq data, termed ChIPWig, which may be executed in a lossless and lossy mode. The algorithm employs delta encoding, run-length encoding and arithmetic encoding akin to smallWig, but in a fundamentally different manner. In particular, ChIPWig aims to smooth out the variance in the average read densities and hence performs a transform step on the relevant data track. Furthermore, it uses specialized coding methods for the location sequence. Unlike smallWig, ChIPWig offers a lossy compression feature through uniform and nonuniform quantization. Uniform quantization ensures best compression results, as it significantly smoothes out the data which may then be efficiently compressed using runlength coding. Unfortunately, and as expected, uniform quantization degrades peak calling accuracy due to the same described smoothing properties. Nonuniform quantization leads to slightly larger file sizes, but does not change the output of peak calling, and hence, motif finding algorithms as it performs smoothing in a fashion controlled by the data distribution.

To illustrate the utility of our lossless and lossy compression methods, we tested the lossless and lossy mode of the algorithm on numerous ChIP-seq files from the ENCODE project. We include detailed information on 10 ChIP-seq files from the ENCODE project and compared the lossless ChIPWig compression rate to those of bigWig, cWig, gzip and the original Wig files. Our results show that lossless ChIPWig compression rates, on average, outperform those of bigWig by 6-fold, gzip 4.5-fold and cWig 2-fold. Lossless ChIPWig also enables random query of different chromosome regions by partitioning the data tracks into blocks of suitable size. In the random access mode, compression is performed on each block individually, trading-off compression rate for ease of targeted access. ChIPWig, unlike bigWig and cWig, also accepts different block sizes. Furthermore, it allows the user to access the summary statistics information (maximum, minimum, average and standard deviation of the average read densities, and peak magnitudes) for each block. Lossy compression is accomplished by first performing uniform or nonuniform quantization of the average read densities, and then executing the lossless ChIPWig procedure. Lossy ChIPWig was tested on the same 10 ChIP-seq files from the ENCODE project, and led to close to 3-fold reductions in file sizes for the case of uniform quantization, and 2-fold reductions in file sizes for the case of nonuniform quantization. For visualization purposes, the nonuniformly compressed files do not introduce any subjective, visible degradation.

In addition, we did a case study on an example ChIP-seq Wig file containing information about chromosome Y. On this dataset and its uniformly and nonuniformly quantized counterparts, we performed peak calling via NarrowPeaks (Madrigal (2016)) (although peak calling is mostly performed on BAM or SAM files, some peak calling methods such as NarrowPeaks allow one to directly operate on Wig files; note that despite its name, NarrowPeaks can identify both narrow and broad peaks). We fed the output of the peak caller into MEME-ChIP (Machanick and Bailey (2011)) for the purpose of identifying binding motif sequences. The results reveal that in contrast to the uniform quantizer, the near-optimal (with respect to mean-square distortion, and for a sufficiently large number of thresholds) nonuniform quantizer did not introduce any performance loss in peak calling and motif finding. In other words, the results produced by NarrowPeaks and MEME-ChIP were identical for the original and nonuniformly quantized files in terms of preserving peak positions and motifs.

The paper is organized as follows. In Section 2, we outline the ChIPWig compression architecture and describe how a ChIP-seq Wig file is processed via delta, run-length, and arithmetic encoding. In Section 3, we first describe the implementation of the lossless ChIPWig algorithm and illustrate its performance with respect to compression/decompression rate and execution time. Then, we proceed to introduce the quantization schemes used and the implementation of the lossy ChIPWig algorithm. In Section 4, we describe the results of peak calling and motif analysis of the example unquantized and quantized ChIP-seq dataset. In Section 5, we discuss the features of the algorithm and Section 6 concludes the paper.

## 2 Methods

### 2.1 Lossless Encoding

In what follows, we outline the initial ChIP-seq Wig file processing via delta and run-length encoding and then proceed to describe the compression technique based on arithmetic encoding. The ChIP-seq Wig files used in our experiments were generated as part of the ENCODE project, where the ChIP-seq files used for visualization are originally stored in bigWig format. We first converted the ChIP-seq files stored in bigWig to Wig format and then performed ChIPWig compression.

A sequence is defined with a capital letter and elements of the sequence are denoted by lower case letters. For instance, *A* = (*a*_1_, *a*_2_, …, *a*_*N*_) is a sequence with *N* elements. We write [*j*] = {1, 2, …, *j*}, for any positive integer *j* ∈ ℕ and [*j, k*] = {*j, j* + 1, …, *k*} for any non-negative integers *j* and *k* where *j* < *k*.

The ChIP-seq Wig files comprise the following sequence tracks:

- *The Location Sequence L* = (*ℓ*_1_, *ℓ*_2_, …*ℓ*_*N*_), where *N* denotes the length of the sequence and *ℓ*_*i*_ ∈ ℕ for all *i* ∈ [*N*] satisfies *ℓ*_*i*_ < *ℓ*_*i*+1_, *i* ∈ [*N* − 1]. In this sequence, one often has *ℓ*_*i*+1_ = *ℓ*_*i*_+*s*, where *s* > 1, and *i* ∈ [*N*], except for some skipped locations, for which *ℓ*_*i*+1_ > *ℓ*_*i*_ + *s*. We note that while in most of the tested ChIP-seq data, *s* is a constant and is equal to 25, in some cases *s* = 21 and *s* = 10 were used as well. We also observed that locations for which *a*_*i*+1_ > *a*_*i*_ + *s* are the exception rather than norm, but there is still a significant number of them. As an example, in one prototypical ChIP-seq Wig file used as the running example and described in the Results section, the number of locations for which *s* > 25 equals 6,723,719, while the number of locations for which *s* = 25 equals to 73,575,541.
- *The Average Read Density Sequence C* = (*c*_1_, …, *c*_*N*_), where *c*_*i*_ ∈ ℝ^+^ for all *i* ∈ [*N*]. The *c*_*i*_’s indicate the average read density corresponding to the location *ℓ*_*i*_. The sequences *L* and *C* have the same length.

Having defined the sequences *L* and *C* which represent the two columns of an ENCODE ChIP-seq Wig file, we are ready to describe the lossless compression techniques applied on the sequences.

Compressing the Location Sequence *L* involves the following steps.

1. **Delta encoding:** Refers to computing the Difference Sequence *D* = (*d*_1_, *d*_2_, …, *d_N_*), *d_i_* ∈ ℤ, *i* ∈ [*N*], of a sequence. When the underlying sequence equals *L*, one has *d_i_* = *ℓ*_*i*_ − *ℓ*_*i*−1_.
2. **Run-length encoding:** Results in two sequences *S* = (*s*_1_, …, *s_K_*), *R* = (*r*_1_, …, *r_K_*) obtained by performing run-length encoding on the delta encoded sequence *D*, where *s_i_* ∈ *D* captures the symbol, while *r_i_* equals the number of consecutive appearances (i.e., the run-length) of *s_i_* in *B*, for all *i* ∈ [*K*].
3. **Arithmetic encoding:** Is a form of entropy coding that uses the symbol probability distribution to perform variable-length interval parsing that leads to a sequence being represented by an interval. Arithmetic coding often outperforms Huffman entropy coding, as it operates on the sequence as a whole, rather than on individual symbols. The sequences *S* and *R* are compressed by an arithmetic encoder.

The compression method for the Average Read Density Sequence *C* involves the following steps.

1. **Scaling:** Refers to converting the Average Read Density Sequence, which consists of non-negative real-valued numbers, into integers. Each *c_i_* ∈ *C* is multiplied by a scaling factor 10^*E*^, for some *E* ∈ ℕ, where *E* is chosen based on the number of decimals in *c_i_*. This results in a sequence *A* = (*a*_1_, …, *a*_*N*_), *a*_*i*_ ∈ ℕ, *i* ∈ [*N*], where the *a_i_*’s are integer-valued. In most ENCODE ChIP-seq Wig files, the average read densities are represented with at most four decimal digits, resulting in *E* = 4. In lossless ChIPWig, all four decimal digits are retained, while in the lossy implementation, only up to two decimal digits are preserved.
2. **Delta encoding:** As before, this form of coding results in a Difference Sequence *W* = (*w*_1_, …, *w_N_*), *w_i_* ∈ ℤ, *i* ∈ [*N*] is generated according to *w_i_* = *a_i_* − *a*_*i*−1_.
3. **Arithmetic encoding:** Is applied on the delta encoded sequence *W*, separately from *S* and *R*. Joint compression is not pursued, as extensive analysis of joint track statistics performed for smallWig (Wang *et al*. (2016)) reveals that joint encoding for Wig files in general does not significantly improve performance, but increases computational time.

The block diagram of the compression scheme for the Location Sequence and the Average Read Density Sequence is shown in Figure 1(a). The lossless ChIPWig performs scaling, delta encoding and arithmetic encoding on the average read density sequence, while the lossy ChIPWig adds a quantization step to the Read Density sequence. The quantized data is then processed by scaling, delta encoding and arithmetic encoding. The decompression method of ChIPWig is illustrated in Figure 1(b). Both the lossless and lossy ChIPWig perform the same decompression steps.

**Figure 1:**
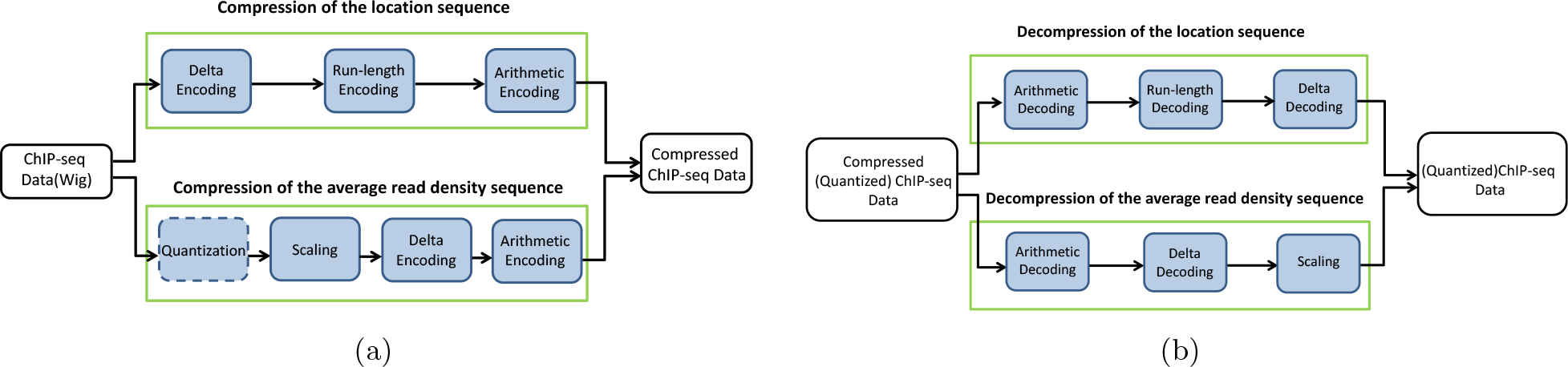
The block diagram of the ChIPWig compression and decompression algorithms. a) compression model; b) decompression model.

### 2.2 Quantization

The output of a ChIP-seq experiment is sequence information in FASTQ format which is used for downstream analysis that generates a vast amount of other types of data. A typical pipeline includes read mapping, peak calling, and motif discovery, which in turn produce SAM/BAM, Wig, and Bed files. There is a storage and computation time trade-off for each of the file types: keeping all files incurs a high cost of storage, while re-running large scale analysis is computationally expensive. A solution is to store raw data in a lossless manner, and highly processed data in a lossy form. In particular, Wig files contain highly “processed” counts for protein binding sites, often containing read counts themselves, averaged over a window of 25 bp, or fold-control ratios. One advantage of the Wig files is a simple data structure that can accommodate visualization on genome browsers, which does not require large precision due to low sensitivity of the human eye. For these reasons, we suggest lossy compression of read densities site information in Wig files using scalar quantization schemes that map the large set of average read densities into a significantly smaller one. We investigate both uniform and nonuniform scalar quantization.

The exposition in this section is heavily based on definitions, concepts and formulas from classical sources on quantization (Gallager (2006), Gersho and Gray (1992)). Details regarding density estimation and relevant quantization results are deferred to the Supplementary Material.

#### Scalar Quantization

A scalar quantizer divides the set of real numbers or a given interval into *M* subsets (subintervals) *Q*_1_, …, *Q_M_*, also called quantization regions. Each quantization region *Q_i_* = (*t_i_, t*_*i*+1_) contains a chosen representative point *ω_i_* ∈ *Q_i_*, termed a level. Quantization refers to mapping values in *Q_i_* to the level *ω_i_*, for all *i* ∈ [*M*]. The set {*t*_0_, *t*_1_, …, *t_M_* } in which (*t_i_, t*_*i*+1_) defines the interval *Q_i_*, is called the set of thresholds and the set {*ω*_0_, *ω*_1_, …, *ω*_*M*−1_} is called the set of levels. Thus, a scalar quantizer may be viewed as a function *Q*(*x*): ℝ → ℝ where *Q*(*x*) = *ω_i_* if *x* ∈ *R_i_* (Gallager (2006)).

Let *X* = (*X*_1_, *X*_2_, …, *X_N_*) be a sequence of i.i.d random variables with individual probability density function *f_X_i__* (*x*) or some adequate distribution function, whenever the variables are discrete. Assume that the variables are quantized to *Y* = (*Y*_1_, *Y*_2_, …, *Y*_*M*_) and let *Q*(*X*) be the quantizer function used in the process. Then, the *second order distortion* is defined as follows:

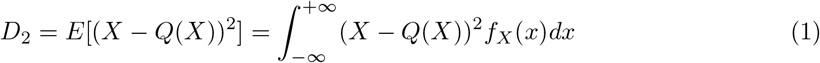

Since the second order distortion measures the difference between the original data ad quantized data, the goal is to design a quantizer function that minimizes *D*_2_, given *X* and *f*_*X*_ (*x*). The second order distortion is also called the mean-squared distortion or the mean-squared error (MSE). This distortion may be generalized for arbitrary orders *m, m* ≥ 1, and the choice of the distortion function may be governed by the downstream processing tasks performed on the quantized data. For example, if one seeks very accurate quantization for large values, a distortion with large value of *m* may be desirable. However, we only consider the case that *m* = 2, as this choice is adequate for our implementation and describe optimal quantizer derivations for *m* ≥ 2 in the Supplementary Material.

We make use of both uniform and nonuniform quantizers and state closed-form formulas the MSE in both cases, which are known from the quantization literature. The formulas are applied to the estimated data distributions as outlined in the Supplementary Material.

#### A. Uniform Scalar Quantization

In uniform scalar quantization, each quantization level *Q_i_* has the same length *|Q_i_|* = Δ, provided that the underlying distribution has finite support. Each value *x* ∈ *Q_i_* = (*t_i_, t*_*i*+1_) is mapped to the midpoint 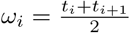. The uniform quantization function takes the form:

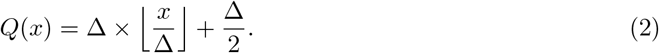

An example of a uniform quantizer with *M* = 7 quantization levels is shown in Figure 2. When the data is supported on an infinite interval, one of the quantization intervals has to be of infinite length, and the corresponding level has to be chosen with care. Given that all our data is supported in a finite interval, we do not discuss this scenario in detail.

**Figure 2:**
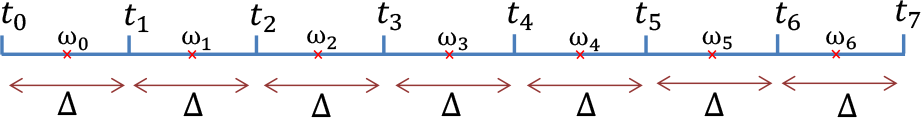
An example of a uniform quantizer with seven quantization levels, where 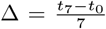 and 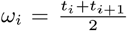 for *i* ∈ [0, 6].

It is known that for a uniform quantizer with *M* quantization levels between *a* and *b* and 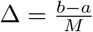, one has

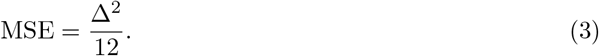

The proof for the general *m*-th order distortion result is retraced in the Supplementary Material.

#### B. Nonuniform Scalar Quantization

In nonuniform quantization, the lengths of quantization regions are not necessarily equal. Each value *x* ∈ *Q_i_* = (*t_i_, t*_*i*+1_) is mapped to either 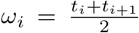, or another carefully selected level. Here, we only pursue the former level implementation. An example of a nonuniform quantizer with seven quantization levels is given in Figure 3, where the Δ_*i*_’s, for *i* ∈ [1, 7], are the length of the quantization regions.

**Figure 3:**
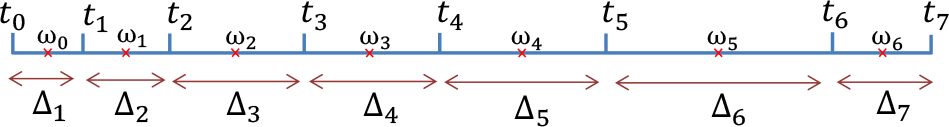
An example of a nonuniform quantizer with seven quantization levels, where Δ_*i*_ = *t*_*i*_ − *t*_*i*−1_ for *i* ∈ [1, 7] and 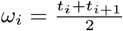 for *i* ∈ [0, 6].

To design a quantization scheme with *M* quantization levels, when *M* is large, one usually refers to *asymptotic quantization theory* (Gallager (2006). The *point density* of a quantizer is defined as a function *λ*(*x*) such that for any *a* and *b, a* < *b*,

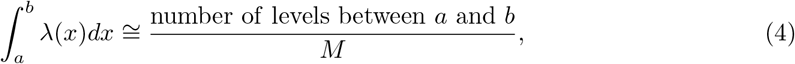
where *λ*(*x*) ≥ 0 and 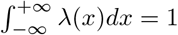.

It can be shown that for a nonuniform quantizer with *M* quantization levels, the MSE depends on the probability density function *f*_*X*_ (*x*) and the point density function *λ*(*x*) as:

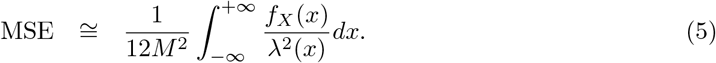

The point density that minimizes the MSE reads as

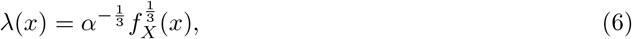
where the constant *α* is obtained from 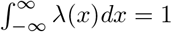.

The extension of this result for the general *m*-th order distortion is outlined in the Supplementary Material.

Thus, given *f*_*X*_ (*x*), the constant *α* and the density *λ*(*x*) may be obtained from 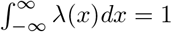 and (6), respectively. Thus, given the number of quantization levels *M*, using (4), the number of quantization levels between any two values in the support of the distribution, *υ*_1_ and *υ*_2_, *υ*_1_ < *υ*_2_ can be obtained. Hence, asymptotically optimal (with respect to the MSE and for a large number of levels) quantizer design may be guided by the point density, as we illustrate on ChIP-seq Wig file data in the Supplementary Material.

## 3 Results

We start by describing the features of the lossless ChIPWig method both in the standard mode and random query mode and present a set of results pertaining to compression rates, compression and decompression times for 10 test files downloaded from the ENCODE hg19 browser. We then proceed to present the same result for the lossy mode, which we accompany with a case study of downstream peak calling and motif finding on quantized data.

### 3.1 Lossless ChIPWig

As explained in Section 2, we use separate arithmetic encoders/decoders to process the sequences *S, R* and *W*. The arithmetic encoding method that is used for our implementation is the same as the one used in smallWig (Wang *et al*. (2016)). This arithmetic compression algorithm is based on range coding (Martin (1979)) and some techniques from the rangemapper by Polar (http://ezcodesample.com/reanatomy.html?Source=To+article+and+source+code). For more details, the reader is referred to (Wang *et al*. (2016)).

The random access function is implemented through *block-wise* encoding, in which a ChIP-seq Wig file is divided into blocks of fixed length *b* where *b* = 2^*k*^, *k* = 11, ⋯, 18, and where each block is processed and encoded separately. The encoded blocks are merged into one file which forms the compressed file. When the random query mode is active, a summary statistics of each block is stored in the compressed file. The summary statistics includes: (i) the minimum average read density in the block, (ii) the maximum average read density in the block, (iii) the mean of average read densities in the block and (iv) the standard deviation of the average read densities in the block. We also store two more values for each block that are not usually recorded in Wig files. These values represent the minimum jump and maximum jump in read density, which are of relevance for identifying relevant peak regions, and are defined as follows. Each local maxima in the average read density is identified with its value and the average densities appearing before and after the maxima. A jump is defined as the difference between the maxima and the average read densities immediately preceding and following the maxima. The minimum jump and the maximum jump are the smallest and largest jump within each individual block. For our experiments, we used 10 ChIP-seq data files in bigWig format from the ENCODE hg19 browser listed in Table 1 which were converted to Wig format.

**Table 1:** Names and sizes (in MB) of the 10 ChIP-Seq files used in the compression experiments, retrieved from the ENCODE hg19 browser.

| File Number | File Size | File Name |
| --- | --- | --- |
| 1 | 959 | wgEncodeBroadHistoneA549H3k09acEtoh02Sig |
| 2 | 1000 | wgEncodeBroadHistoneA549CtcfEtoh02Sig |
| 3 | 938 | wgEncodeBroadHistoneA549CtcfDex100nmSig |
| 4 | 1460 | wgEncodeBroadHistoneA549ControlEtoh02Sig |
| 5 | 1210 | wgEncodeBroadHistoneA549H3k04me3Etoh02Sig |
| 6 | 925 | wgEncodeBroadHistoneA549H3k27acEtoh02Sig |
| 7 | 1090 | wgEncodeBroadHistoneA549H3k27me3Dex100nmSig |
| 8 | 781 | wgEncodeBroadHistoneA549H3k79me2Dex100nmSig |
| 9 | 1020 | wgEncodeBroadHistoneCd20ro01794Ezh239875Sig |
| 10 | 1210 | wgEncodeBroadHistoneDnd41Ezh239875Sig |

Figure 4 shows the compression rates achieved by the ChIPWig, compared to the rates of gzip and bigWig (The compression rate equals the ratio of the compressed file size and the uncompressed file size). ChIPWig offers 6-fold rate improvement compared to bigWig and almost 4.5-fold improvement compared to gzip. For compression with random queries with block size 65536, ChIPWig offers a 5.5-fold rate improvements compared to bigWig. In order to compare the size of the compressed files with ChIPWig and cWig, we compared the average size of the ChIP-seq Wig files that we obtained with those reported in the paper introducing cWig, and found that our method offers almost 2-fold decrease in file size compared to cWig. We had to resort to this type of comparison as cWig is no longer available online.

**Figure 4:**
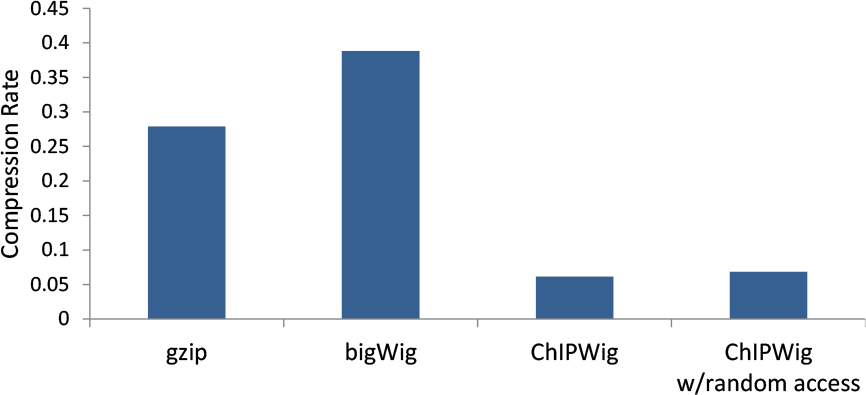
Compression rates achieved by gzip, bigWig, and ChIPWig algorithms both in the standard mode and the random query mode with block size 65536. Results represent the average over 10 sample ENCODE files.

In Figure 5, we present the running time of the ChIPWig compression/decompression schemes, as well as those of gzip and bigWig. In general, the running time of ChIPWig is slightly longer, but comparable, to that of bigWig and gzip. The average compression times in the standard mode of ChIPWig is on average 0.12 seconds longer than that of in bigWig and gzip, while the average decompression time is 0.15 and 0.17 seconds longer than that of in bigWig and gzip, respectively. To compare the effect of different block sizes used for random query on compression rate and compression/decompression time, we refer the reader to Figure 6. In these experiments, the block sizes ranged from 4096 to 65536. In general, the results show that as the size of the block increases, the compression/decompression time and compression rate decrease. The ChIPWig algorithm with random access leads to a slight increase in compression rate from 0.06114 in standard mode to 0.1067 in random access mode with the block size 4096. It also leads to a modest increase from 0.2066 to 0.3443 MB/sec in the compression time rate and an increase from 0.1684 to 0.2252 MB/sec for the decompression time rate when compared to the standard mode. We also ran the ChIPWig algorithm in random query mode to find the average decompression time of a query of length 1000 in a compressed ChIP-seq file which was compressed by ChIPWig using blocks of size 65536. The results show that the average query time for queries of length 1000 over all 10 Wig files and 10 queries on each file are close to 0.24 seconds. This decompression time is longer than that of bigWig, which roughly amounts to 0.003 seconds. ChIPWig also provides a statistical analysis during queries and returns the minimum, maximum, average and standard deviation of average read densities.

**Figure 5:**
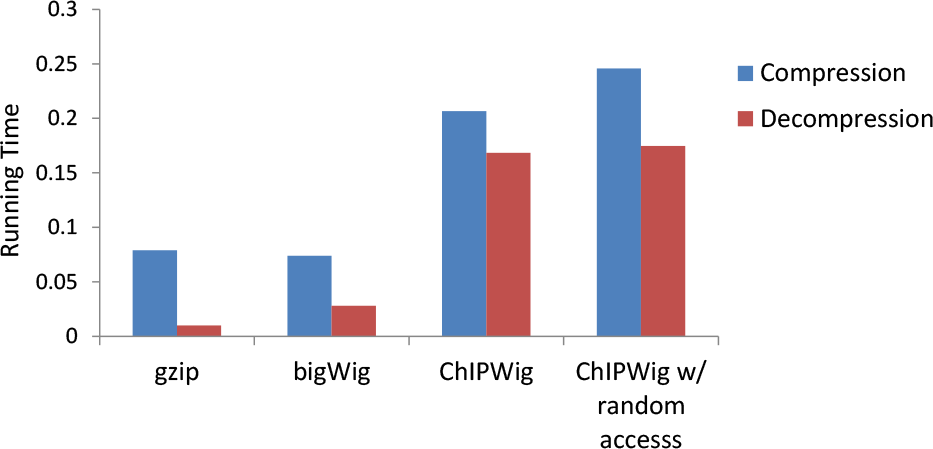
Compression and decompression times of the gzip, bigWig, and ChIPWig algorithms in standard mode and the random query mode with block size 65536. The compression and decompression times are expressed in seconds per MB of the original Wig file size. All the results were averaged over the example 10 sample files from ENCODE hg19.

**Figure 6:**
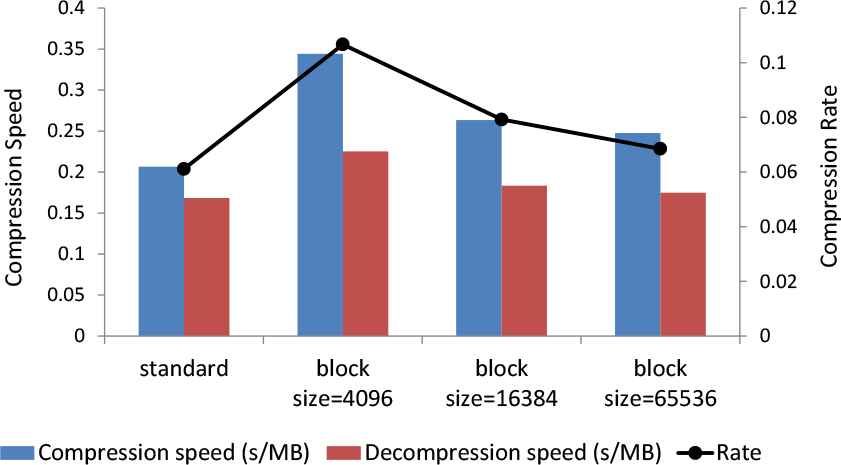
Compression rate, compression time, and decompression time for different block sizes. The label “standard” indicates that the whole sequence is compressed as a single block. The compression/decompression time is expressed in seconds per MB in the original file. The *y*-label is used for both the rate and the speed (s/MB). All the results were obtained by averaging over the selected 10 sample ENCODE files.

In our second round of tests, we also applied both nonuniform and uniform quantization schemes on the 10 tested ChIP-seq Wig files. We first set a threshold *τ* on the average read density and applied the designed quantization schemes on values that lie on (0, *τ*). For values outside of this interval, we used one level as specified in the Supplementary Materials. For our analysis, we first assumed that *τ* = 50 and designed quantizers with *M* = 50 levels. Then, we repeated the simulations by letting *τ* be 70 percent of largest average read density in the file and let *M* = 50 and *M* = 100. For nonuniform quantization, we used an optimized point density for these two choices of the cutoff threshold *τ*.

The average ChIPWig compression rate of the quantized files is shown in Figure 7. Overall, the uniform quantizer has a lower compression rate than the nonuniform quantizer with the same number of quantization levels and threshold. While the average compression rate in the lossless standard mode is almost 0.06, with nonuniform quantization, the average compression rate is reduced to 0.042 when *M* = 50 and *τ* = 50; for uniform quantization, it is reduced to 0.024 for the same parameters. For the nonuniform quantizer, there is a small increase in the average compression rate from 0.023 for *M* = 50 and *τ* equals to the 70 percent of the maximum average read density in the file to 0.042 for *M* = 50 and *τ* = 50. The results for the uniform quantizer also shows an increase in the average compression compression rate from 0.006 to 0.024, where the former result is obtained for *M* = 50 and *τ* equal to the 70 percent of the maximum average read density density in the file; the latter is obtained for *M* = 50 and *τ* = 50.

**Figure 7:**
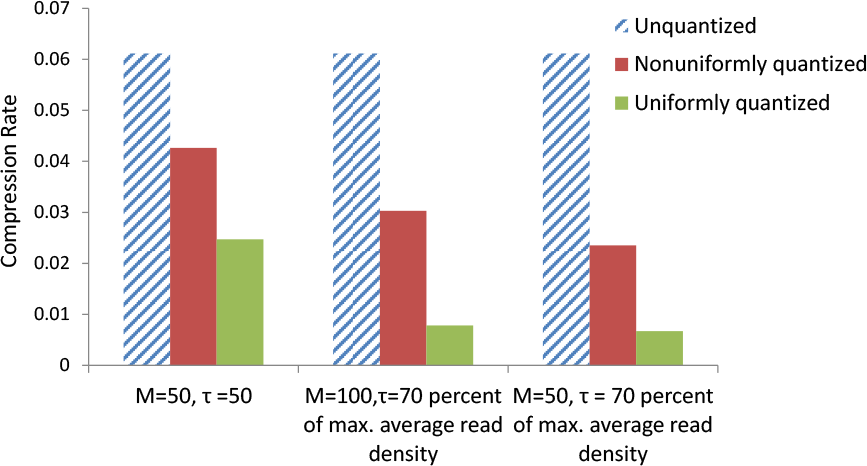
Compression rates of the 10 sample ChIP-seq Wig files before and after nonuniform and uniform quantization schemes with *M* = 100 and *M* = 50. The threshold *τ* is set to 50 or 70 percent of the maximum average read density in each of the files.

## 4 Peak Calling and Motif Analysis

### 4.1 Peak calling

One of the most critical steps in ChIP-seq data analysis is to identify enriched regions and discover potential binding sites of proteins. Generally, a peak is called when the number of covering reads exceeds a predetermined threshold, or when the enriched region is statistically significantly compared to the background peaks (Bailey *et al*. (2013), Nakato and Shirahige (2016), Steinhauser *et al*. (2016)). Among frequently used peak calling algorithms are MACS (Zhang *et al*. (2008)), SPP (Kharchenko *et al*. (2008)) and NarrowPeaks (Mateos *et al*. (2015)). NarrowPeaks operates on ChIP-seq data in Wig and bigWig formats (Mateos *et al*. (2015)) and despite its name, recovers both narrow and broad peaks. We hence focus our attention on analyzing the performance of this peak caller on lossless and quantized Wig files.

To see how quantization affects peak calling, we use the first file in Table 1, “wgEncodeBroad-HistoneA549H3k09acEtoh02Sig” as our running example and focus on the locations and average read densities reported for chromosome Y; for simplicity, we refer to this file as the chrY ChIP-seq file. NarrowPeaks is applied with the default parameters both on the chrY ChIP-seq file and the corresponding (non) uniformly quantized files.

NarrowPeaks returns the list of peaks with the start position, end position, and width of the peak, as well as the peak position and its score. As an illustration, the first five peaks of the chrY ChIP-seq file identified by NarrowPeaks are listed in Table 2.

**Table 2:** The first five peaks identified by the NarrowPeaks when applied to chrY ChIP-seq file. The symbol “*” denotes that the strand information is not used.

| Index | Chr. | Start | End | Width | Strand | Peak Position | Score |
| --- | --- | --- | --- | --- | --- | --- | --- |
| 1 | ChrY | 2649426 | 2649625 | 200 | * | 2649426 | 2 |
| 2 | ChrY | 2649701 | 2650500 | 800 | * | 2650051 | 12 |
| 3 | ChrY | 2650676 | 2651150 | 475 | * | 2650751 | 3 |
| 4 | ChrY | 2653751 | 2653975 | 225 | * | 2653776 | 3 |
| 5 | ChrY | 2654326 | 2654500 | 175 | * | 2654326 | 2 |

The compression rates of the uniformly and nonuniformly quantized chrY ChIP-seq file are shown in Figure 8. As expected, the uniform quantizer has a lower compression rate compared to the nonuniform quantizer. Both uniform and nonuniform quantizer have the lowest compression rate when *M* = 50 and *τ* = 307. The compression rate for the nonuniform quantizer shows a slight increase from 0.032 for *M* = 50, *τ* = 307 to 0.045 for *M* = 50, *τ* = 50 and the compression rate for the uniform quantizer has also an increase from 0.012 to 0.025 for the same parameters as the nonuniform quantizer.

**Figure 8:**
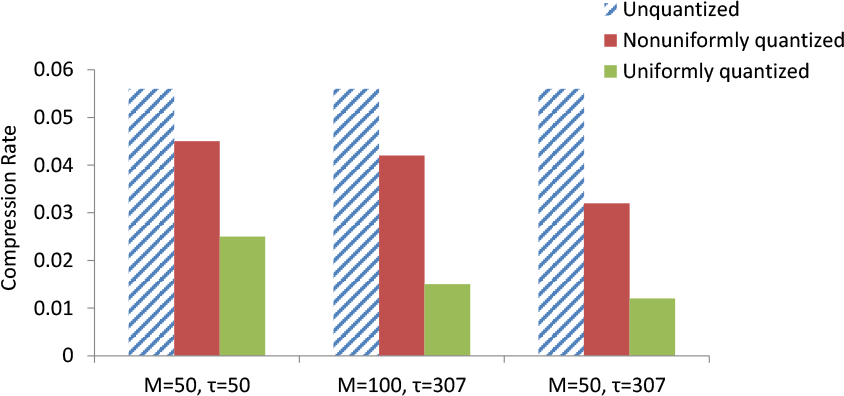
Compression rates of the chrY ChIP-seq file before and after performing uniform and nonuniform quantizations. The portion of the file allocated to chromosome Y is 5 MB.

To evaluate the accuracy of the peak positions and scores found based on the quantized files, we calculated the correlation coefficient between the value found for the unquantized file and the quantized file with *M* = 50 and *τ* = 50. Details of the derivations may be found in the Supplementary Material. We found that the correlation coefficient of the peak positions and scores between the unquantized file and the nonuniformly and uniformly quantized file are 1 and 0.8663, respectively. The correlation coefficients show that there is a perfect linear relationship in the peak positions of the unquantized and nonuniformly quantized files and that consequently, nonuniformly quantized files may replace nonquantized files without causing peak caller performance degradation.

### 4.2 Motif analysis

A widely accepted assumption is that many transcription factors exhibit DNA binding motifs so as to successfully locate themselves in the genome. Thus, discovering DNA motifs from the peaks is another step in the downstream analysis of ChIP-seq data. Although approaches may vary by software packages, the general workflow involves finding overrepresented sequences among a set of sequences covering the peak positions (Park (2009)). MEME (Bailey *et al*. (2006)) is the most popular tool for accomplishing this task, and it is based on the Expectation-Maximization (EM) algorithm. The authors of MEME have also developed MEME-ChIP (Machanick and Bailey (2011)) to accommodate large datasets. To demonstrate that quantization has little or no effects on the motif downstream analysis, we discovered motifs from the peaks identified in the previous testing stage.

For each of the seven generated peak files (one for the original, unquantized file, three for the uniformly and three for the nonuniformly quantized file), we ran MEME-ChIP in default mode as described in what follows. We first computed the center of the peak regions by taking the floor(median(start,end)), and extended the sequence by 150 base pairs in both directions. Using the hg19 reference genome, we fetched fasta files containing sequences of 301 bps and ran MEME-ChIP under default setting. Figure 9 shows the three most significant (sorted by E-values) motif logos discovered by MEME-ChIP for the unquantized file. Of our six quantized files, three nonuniformly quantized files discovered the exact same motifs as the original file, verifying that near-optimal nonuniform quantization has no effects on downstream analysis. The remaining uniformly quantized files led to the discovery of similar, but different motifs. We listed all motifs of the files obtained from uniform quantization in the Supplementary Material.

**Figure 9:**
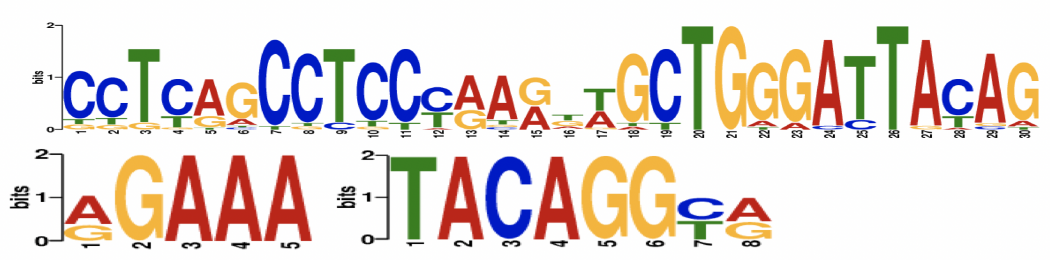
The most significant motif logos discovered by MEME-ChIP for the original unquantized chrY ChIP-seq file and all three nonuniform quantized files.

## 5 Discussion

The basic design principles supporting ChIPWig compression can be easily modified and used on any other Wig data file. In case that ChIPWig is used to compress RNA-seq Wig files with integer expression values, performing run-length encoding on gene expression values is useful to reduce the size of the compressed file, while scaling is redundant and is to be skipped. In the random access mode, ChIPWig operates similarly to smallWig, and uses simple block-based encoding. ChIPWig accepts a variety of block sizes for encoding which is not the case for bigWig and cWig that each encode fixed blocks of size 512. ChIPWig is also implemented both in the lossless and lossy compression modes, while the smallWig, bigWig, gzip and cWig have been implemented only for the lossless compression. In the Supplementary Material, several examples of nonuniformly quantized ChIP tracks are magnified and visualized alongside uncompressed tracks, showing that there is almost no shape distortion which may be of importance for visualization applications. Furthermore, given that nonuniform quantization techniques may be easily designed for any subjective distortion measure and that they tend not to influence peak calling and motif finding in any significant manner, it appears desirable to store these files only in quantized form.

## 6 Conclusion

We proposed a new algorithm termed ChIPWig for compression of ChIP-seq Wig files, operating both in a lossless and lossy mode. ChIPWig has a significantly lower compression rate compared to bigWig, gzip and cWig. ChIPWig also has random access feature and provides access to summary statistics of each block. The performance of lossless and lossy ChIPWig on different ChIP-seq Wig files from the ENCODE hg19 browser was evaluated in terms of the compression rate and running time in the standard mode as well as the random access mode. Lossy ChIPWig first performs quantization on the average read density sequence of the ChIP-seq files and then uses the same compression method as lossless ChIPWig. Lossy ChIPWig shows an even lower compression rate compared to the lossless ChIPWig, while still maintaining some important features of ChIP-seq data used in peak calling and motif finding.

## Acknowledgements

This work was supported by the National Science Foundation Emerging Frontiers for Science of Information Center, the National Institute of Health BD2K Targeted software development grant U01 CA198943-02, and the National Science Foundation Graduate Research Fellowship Program [DGE-1144245].

